# msBayesImpute: A Versatile Framework for Addressing Missing Values in Biomedical Mass Spectrometry Proteomics Data

**DOI:** 10.1101/2025.10.02.679746

**Authors:** Jiaojiao He, Barbara Helm, Franziska Gödtel, Katharina Büchner, Marcel Schilling, Marc A. Schneider, Laura V. Klotz, Hauke Winter, Britta Velten, Ursula Klingmüller, Junyan Lu

## Abstract

Advancements in mass spectrometry (MS) technologies have significantly improved the ability to quantify proteins and analyse their modifications. However, MS-based proteomics datasets frequently encounter missing values due to a complex interplay of missing at random (MAR) and missing not at random (MNAR) mechanisms. If unaddressed, such missing data can result in information loss and biased outcomes in data pre-processing, such as normalisation, as well as subsequent analyses and interpretations. Few approaches effectively address both MAR and MNAR, and those that do often necessitate manual tuning of mixture percentages between them or rely on two-group experimental designs. To enhance the handling of missing values, we developed msBayesImpute, an innovative computational method that integrates Bayesian factorization with probabilistic dropout models. We evaluated msBayesImpute against several popular imputation methods using both simulated missing values and those generated through a dilution series experiment on samples from lung cancer patients. Our comprehensive benchmark demonstrated superior performance in reconstructing missing values, estimating normalization factors, and identifying differentially expressed proteins across varying levels of missingness. Notably, msBayesImpute does not require predefined experimental designs and is scalable to large-scale studies. This versatility positions msBayesImpute as an effective and robust tool for enhancing the utility of MS datasets in biological research.

## Introduction

Proteins play a crucial role in the majority of biological processes. Thus, their identification and quantification are essential for understanding physiological and pathological mechanisms, as well as identifying disease biomarkers and therapeutic targets (He and Chiu, 2003; Mikami, Aoki and Kimura, 2012). Proteomics also complements other omics data types, such as genomics, transcriptomics, epigenomics and etc., in studying complex biological systems and diseases (Rinschen and Saez-Rodriguez, 2021; Herbst *et al*., 2022). Recent advancements in mass spectrometry (MS) technologies, especially the label-free acquisition methods, have enabled highly accurate and high-throughput identification and quantification of proteins and their modifications in biological samples (Bantscheff *et al*., 2007; Kumar and Mann, 2009; Patti, Yanes and Siuzdak, 2012). Nevertheless, MS-based proteomics data often suffer from missing values due to the complex interplay of missing at random (MAR) and missing not at random (MNAR) patterns (Lazar *et al*., 2016; Rubin, 1976).

In MS proteomics, MAR values occur independently of protein abundance and primarily due to technical factors such as sample loss, ionization inefficiency, or errors in data processing (Karpievitch, Dabney and Smith, 2012; Wei *et al*., 2018; Finucane, Brennan, and Gormley, 2024). In contrast, MNAR values are abundance-dependent and typically arise when proteins are present near or below the detection limit of instruments (Jin *et al*., 2021). Both patterns often co-exist in datasets, with MNAR being the more prevalent source of missing values (Karpievitch *et al*., 2009). These missing values decrease statistical power and reduce reproducibility across experiments. If not properly handled, the missing values also introduce bias in statistical analysis, such as normalisation, differential expression, and lead to the incorrect interpretation of the data. On the other hand, many popular statistical and machine learning tools, such as principal component analysis (PCA), regularised multiple linear regression, support vector machines (SVM), etc., only work with complete datasets without missing values.

Besides enhancing instrument sensitivity and sample processing, a common approach to addressing missing values MS proteomics data is imputation, which aims to replace missing values with reasonable substitute values. Numerous statistical methods for missing value imputation (MVI) have been developed to date. General-purpose MVI methods, K-nearest neighbours (KNN) (Troyanskaya *et al*., 2001), random forest-based (missForest) (Stekhoven and Bühlmann, 2012), Bayesian PCA (BPCA) (Oba *et al*., 2003), and multi-omics factor analysis (MOFA) (Argelaguet *et al*., 2018) are commonly employed to address datasets containing missing values. These methods reconstruct missing values based solely on observed data, thereby implicitly assuming the MAR mechanism. This assumption can result in biased estimation in MS proteomics data, particularly when the MNAR portion is significant (Jin *et al*., 2021). To manage the MNAR mechanism in MS proteomics data, left-censored imputation methods have been proposed, such as Quantile Regression Imputation of Left-Censored data (QRILC) (Lazar and Burger, 2014), which uses quantile regression to sample from a normal distribution near the lower detection limit, and MinDet (Lazar and Burger, 2014), which replaces missing values with either the dataset’s minimum value or the minimum per sample. However, these methods inherently introduce underestimation errors by disregarding the MAR pattern and generally exhibit inferior imputation accuracy, as they do not leverage the underlying data correlation structure to aid imputation like other MAR imputation methods.

Efforts have been made to model both MAR and MNAR portions in MS proteomics data to enhance imputation accuracy. Gatto and Lilley introduced a mixed imputation method within the R package MSnbase, combining a MAR imputation function, such as KNN, and an MNAR function, such as MinDet (Gatto and Lilley, 2011). However, users must specify the subset of data following the MNAR mechanism, which is typically unknown. Another approach, MsImpute (Hediyeh-Zadeh, Webb and Davis, 2023), can automatically identify protein subsets adhering to MAR or MNAR mechanisms, though its application is confined to two-group experimental designs. Recently, proteomics imputation using self-supervision (PIMMS) (Webel *et al*., 2024) was proposed, aiming to address MS proteomics data imputation with a self-supervised deep-learning framework capable of learning unique missing value patterns inherent in MS proteomics data. Nevertheless, this deep-learning-based method necessitates a considerably large sample size (n>500) to surpass traditional imputation techniques like BPCA, which is often impractical in standard biomedical studies. Given the constraints of the aforementioned methods, there remains a demand for a more flexible and precise MVI method suitable for MS data from typical biological or biomedical studies.

Probabilistic dropout analysis (proDA), which explicitly models the relationship between protein abundance and missing probability using a Bayesian approach, has been shown to enhance differential expression analysis in label-free mass spectrometry data by effectively accounting for mixed MAR/MNAR patterns (Ahlmann-Eltze and Anders, 2020). Drawing inspiration from proDA and Bayesian PCA, here we introduce msBayesImpute, a novel MVI method that integrates probabilistic dropout models into Bayesian matrix factorization to address the unique missing value patterns in label-free MS proteomics data in a data-driven manner.

## Results

### Overview of msBayesImpute

msBayesImpute integrates Bayesian matrix factorization with a probabilistic dropout model to impute missing values in protein abundance matrices derived from label-free mass spectrometry (Figure 1). By combining the strengths of general-purpose methods such as Bayesian PCA (which shares information across features and samples by leveraging covariance structures) with left-censored approaches that address MNAR patterns, msBayesImpute captures both the correlation structure of the data and the abundance-dependent dropout typical of proteomics. Its regularized Bayesian framework suppresses noise while retaining biologically meaningful variation, thereby reducing the risk of false discoveries. Implemented with stochastic variational inference (SVI), the method achieves high computational efficiency and scalability. The detailed description of the architecture and implementation of msBayesImpute can be found in the Methods.

**Figure 1.**
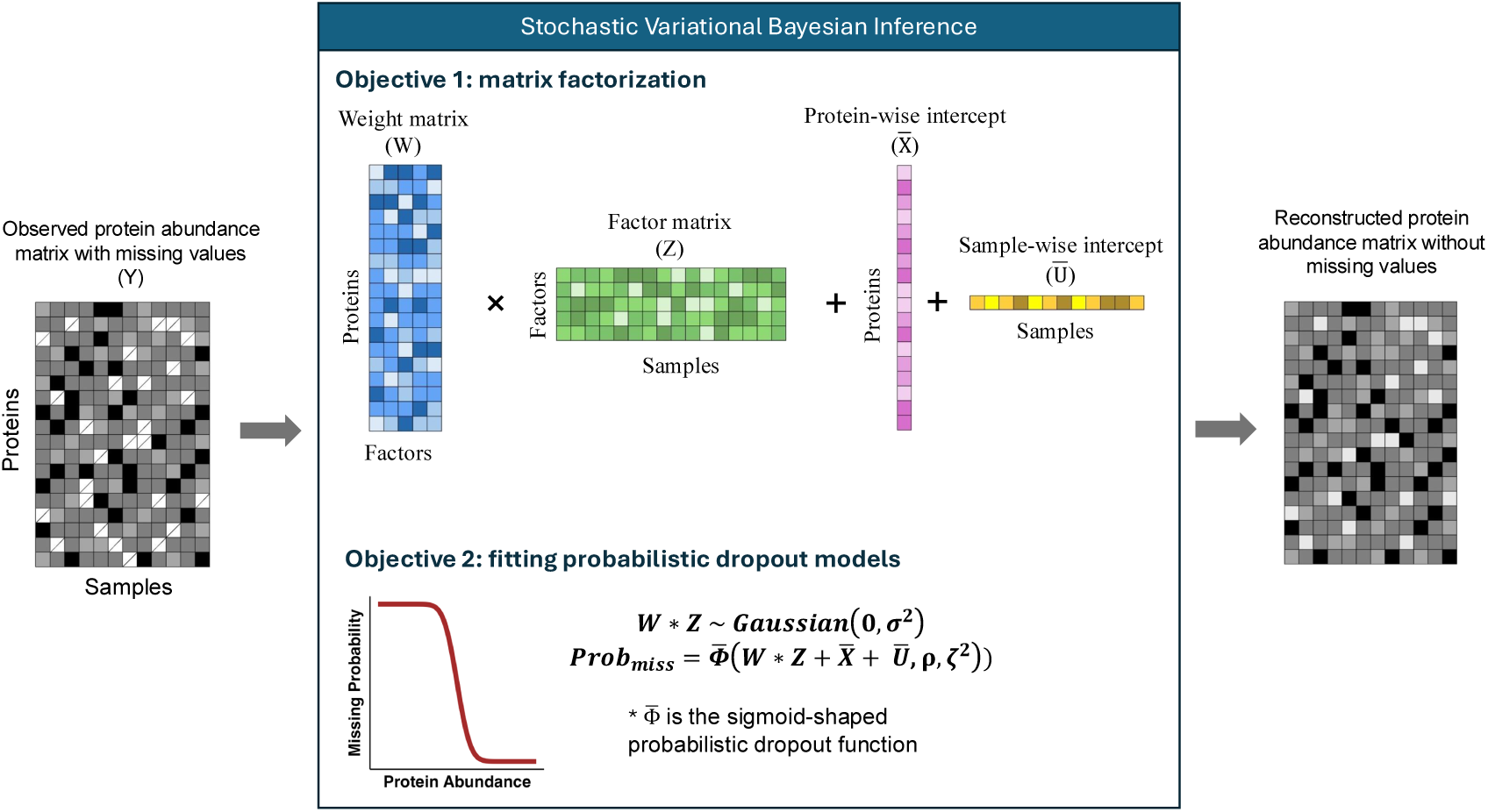
Overview of msBayesImpute. An MS-based proteomics dataset with missing values is decomposed into four components: weight matrix, factor matrix, protein-wise intercept, and sample-wise intercept. msBayesImpute integrates a probabilistic dropout function 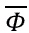 to model abundance-dependent missingness and jointly optimizes these components through stochastic variational inference. The reconstructed matrix without missing values can be directly used in downstream analyses with standard statistical or machine-learning tools.

To facilitate broad adoption, msBayesImpute is available in both R and Python, requires no parameter tuning or predefined experimental design, and can be accessed via a Shiny app or Docker image for non-programmers. This ease of use, combined with its ability to model complex missingness mechanisms, makes msBayesImpute suitable for diverse applications, including exploratory data analysis, normalization, differential expression testing, dynamic modeling of signaling networks, and machine learning tasks. Together, these features position msBayesImpute as a versatile and robust tool for enhancing the reliability and interpretability of label-free MS proteomics in biomedical research.

### msBayesImpute accurately predicts protein-specific dropout functions

In real proteomics datasets, the true dropout functions, which can be visualized as sigmoid-shaped curves describing the relationship between protein abundance and the probability of missingness (Ahlmann-Eltze and Anders, 2020), are unknown, yet accurately estimating them is essential for reliable imputation. Moreover, proteins often exhibit distinct dropout behaviors depending on their physical and chemical properties, further complicating inference when data are limited. To address this, msBayesImpute employs an empirical Bayesian framework to estimate protein-specific dropout functions. In cases where data are sparse—for example, when sample sizes are small or when a protein has too many missing values—the model adaptively shrinks the protein-specific estimates toward global priors derived from all observed data points, thereby ensuring stable and biologically meaningful parameter inference (see Methods).

To directly test the ability of msBayesImpute to recover such dropout functions, we first used purely synthetic data with known parameters. Complete matrices were generated by multiplying Gaussian-distributed weight and factor matrices, representing log-transformed protein abundance data that typically approximate a Gaussian distribution. Each matrix contained 5,000 proteins and 10 latent factors, with a median protein abundance of 20 (see Methods). To assess the effect of sample size, we simulated datasets with 20, 100, and 200 samples. Missing values were then introduced at rates of 10%, 30%, and 50% using a probabilistic dropout function that assigned distinct missingness patterns to proteins. msBayesImpute accurately recovered the shape and parameters of protein-specific dropout functions (inflection point, ρ, and slope, ζ) across all sample sizes and missing rates (Figure 2A–C, Supplementary Figure 1A). Performance improved markedly as sample size increased from 20 to 100, but only modestly from 100 to 200, indicating that msBayesImpute can be effectively trained with relatively small cohorts compared to deep learning–based methods, which typically require hundreds to thousands of samples.

**Figure 2.**
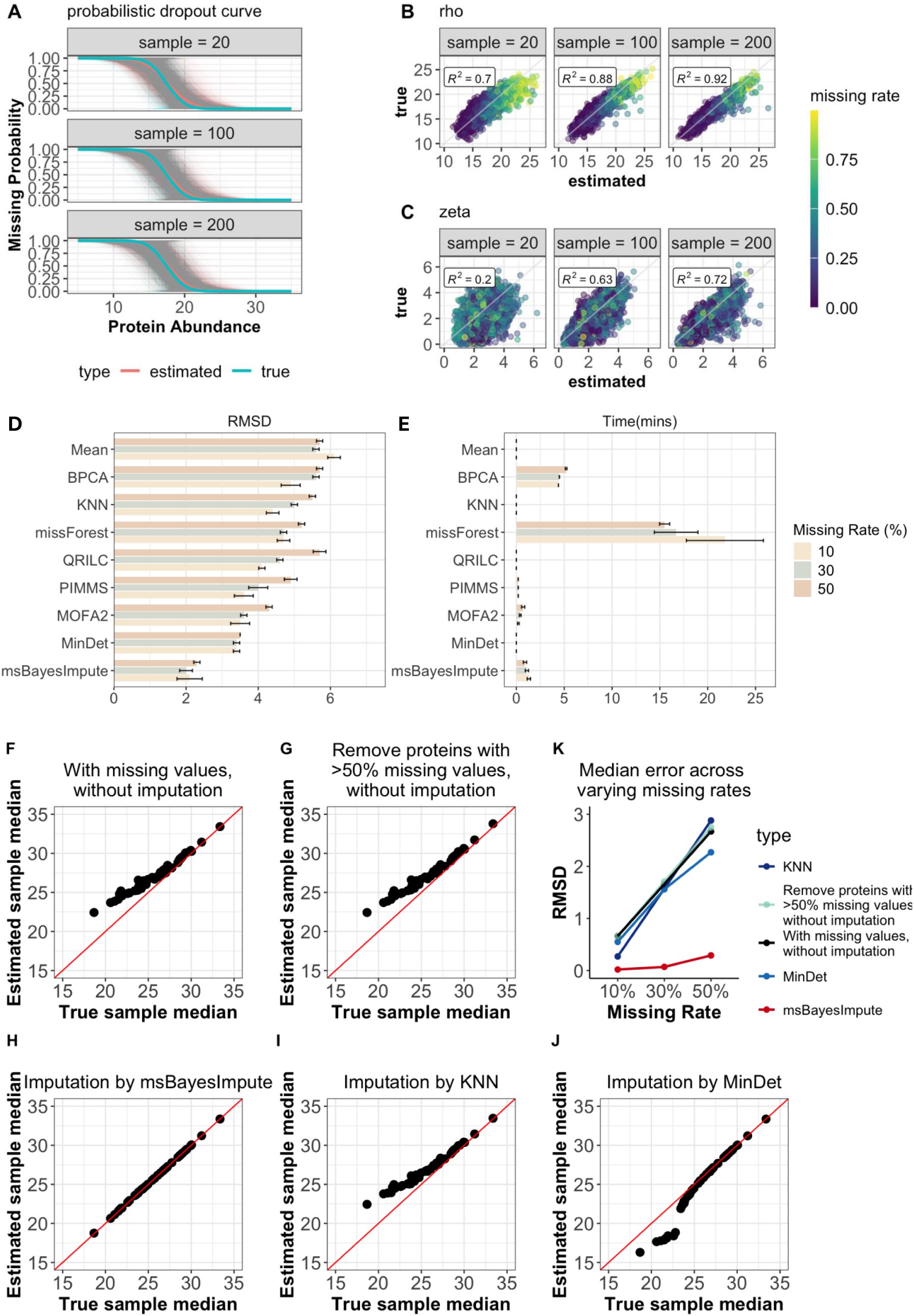
Benchmarking msBayesImpute using synthetic data. (**A)** Comparison of ground truth dropout curves (cyan) and curves inferred by msBayesImpute (red). Solid lines represent the average curve, and shaded lines represent individual protein-specific curves. (**B, C**) Scatter plots comparing true versus estimated dropout parameters (rho and zeta), which define the shape of the dropout curves. R² values indicate the reproducibility of parameter estimation. Each point represents a protein, with color indicating its simulated missing-value probability. (**D)** Root mean square deviation (RMSD) between the imputed and simulated ground truth values. The bars show the averaged RMSD over 5 random repetitions, and the error bars indicate the confidence interval (CI). **(E)** Runtime of each method under the same conditions as in **D**. (**F, G)** Comparison of sample-wise medians derived from ground truth data versus matrices with missing values. In **(G)**, proteins with >50% missingness were excluded before median calculation. **(H–J)** Same analysis as in **F** and **G**, but medians were estimated from imputed complete matrices using msBayesImpute, KNN, and MinDet. **(K)** Median estimation errors across datasets with 10%, 30%, and 50% missing values, comparing msBayesImpute to other approaches.

### Benchmarking the imputation accuracy and computational efficiency with synthetic data

We next benchmarked msBayesImpute against eight commonly used missing value imputation (MVI) methods under varying missing rates (10%, 30%, 50%) in simulated datasets with 50 samples. The compared methods included deterministic minimum (MinDet) (Lazar and Burger, 2014), quantile regression imputation for left-censored data (QRILC) (Lazar and Burger, 2014), Bayesian PCA (BPCA) (Oba *et al*., 2003), multi-omics factor analysis (MOFA) (Argelaguet *et al*., 2018), missForest (Stekhoven and Bühlmann, 2012), k-nearest neighbours (KNN) (Troyanskaya *et al*., 2001), proteomics imputation using self-supervision (PIMMS) (Webel *et al*., 2024), and protein-wise mean imputation. Each condition was repeated on five datasets with the same missingness profile. Among all methods, msBayesImpute showed the best performance, yielding the lowest RMSD and therefore the highest imputation accuracy, with MinDet and MOFA ranking next (Figure 2D). PIMMS performed well at low missingness (10%) but degraded sharply as missingness increased, a trend also observed for KNN, QRILC, and BPCA. In contrast, msBayesImpute, MinDet, missForest, and protein-wise mean imputation displayed relatively stable RMSD across missingness levels. Notably, missForest and BPCA were computationally inefficient for high-dimensional matrices, while msBayesImpute achieved completion in ∼1 minute (Figure 2E).

### msBayesImpute improves normalization accuracy

Most normalization methods used in MS proteomics—such as median normalization or variance stabilizing transformation (VSN) (Huber et al., 2002) —were originally developed for gene expression data, where missing values are rare. These methods assume that the overall distribution of feature values is comparable across samples, with similar medians or means. In proteomics, however, missing not at random (MNAR) values violate this assumption: low-abundance proteins are more likely to drop out, which inflates sample medians and biases the size factors used for normalization. To assess this effect, we simulated datasets with ∼30% missing values. Sample medians derived from incomplete matrices were consistently shifted upward relative to the ground truth (Figure 2F–G). Filtering proteins with >50% missingness did not correct the bias and instead discarded informative data.

We next benchmarked msBayesImpute against representative MAR– and MNAR-based methods (KNN and MinDet, respectively) in recovering true sample median abundances. Although both KNN and MinDet partially reduced the bias, their imputed medians still deviated noticeably from the ground truth (Figure 2H–J). In contrast, msBayesImpute consistently reconstructed medians that closely matched the true values. Across all tested missingness levels (10%, 30%, 50%), it achieved the lowest and most stable normalization error (Figure 2K). Together, these results show that msBayesImpute accurately recovers sample-wise size factors even under substantial dropout, thereby providing a more robust foundation for downstream analyses such as differential expression.

### Benchmarking on semi-synthetic HeLa cell line proteomics data highlights the robust and accurate performance of msBayesImpute

To evaluate msBayesImpute on real-world data and benchmark it fairly against existing methods, we used a published HeLa cell line proteomics dataset that was originally employed to develop and assess PIMMS, a state-of-the-art deep learning–based imputation method (Webel et al., 2024). The dataset consisted of 564 HeLa runs acquired on a Q Executive HF-X Orbitrap instrument during continuous MS quality control. In addition, a smaller dataset with 50 samples was generated to test performance dependence on sample size (Webel et al., 2024). For benchmarking, missing values were introduced using the approach of Lazar et al. (2016), which has been widely applied in proteomics benchmarking studies (Harris *et al*., 2023, Jin *et al*., 2021, Wei *et al*., 2018, Webel et al., 2024). This method provides precise control over both the overall missing rate and the mixture of missing not at random (MNAR) and missing at random (MAR) patterns. In the MNAR setting, low-abundance values are preferentially removed to mimic left-censoring near the detection limit, whereas in the MAR setting, values are masked independently (see Methods). We generated a series of data matrices with MNAR proportions of 0%, 25%, 50%, 75%, and 100% for both the smaller (*N*=50) and larger (*N*=561) datasets after filtering out proteins with greater than 75% missing values and samples with greater than 50% missing values.

Across both small and large datasets, msBayesImpute demonstrated the best reconstruction accuracy and maintained stable performance across all MNAR levels. In contrast, methods based on MAR assumptions—including missForest, MOFA, PIMMS, BPCA, KNN, and mean imputation—showed steadily deteriorating accuracy as the MNAR proportion increased (Figure 3A–B). Methods explicitly modeling MNAR, such as QRILC and MinDet, improved with higher MNAR content but remained substantially less accurate overall.

**Figure 3.**
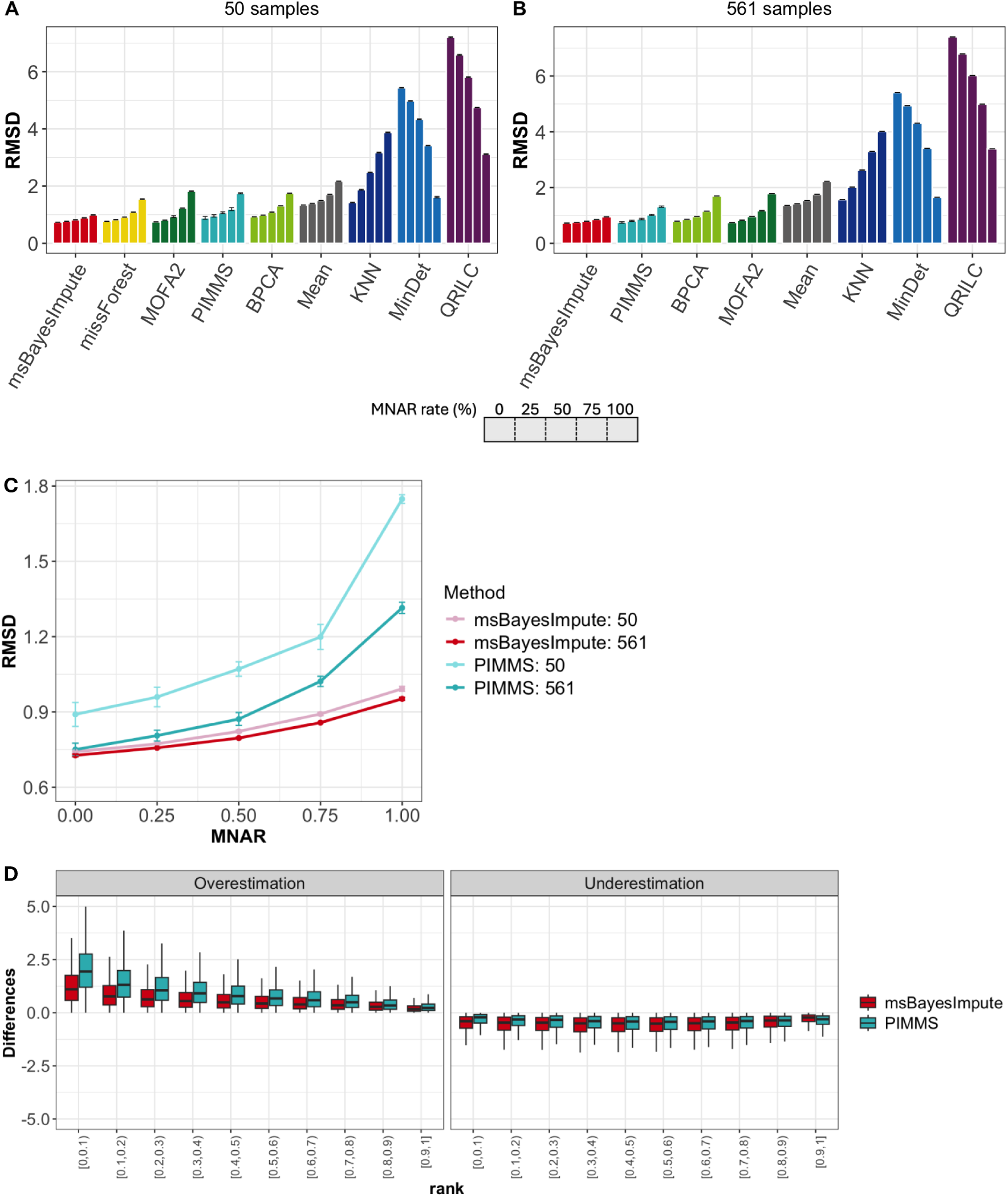
Benchmarking msBayesImpute on semi-synthetic published HeLa cell line proteomics data with varying MNAR proportions. **(A)** Reconstruction accuracy of nine imputation methods across datasets with MNAR proportions ranging from 0% to 100% (25% increments from left to right), based on RMSD between imputed and ground truth values in the smaller dataset (*N*=50). **(B)** Performance of the same methods on the larger dataset (*N*=561), excluding missForest, which failed to converge. **(C)** Direct comparison of msBayesImpute and PIMMS across both small (*N*=50) and large (*N*=561) datasets, showing RMSD as a function of MNAR proportion. **(D)** Protein-level imputation error analysis comparing msBayesImpute and PIMMS, with over– and underestimation quantified across ten ranked protein abundance groups.

In the small dataset (*N*=50), missForest performed second best but was computationally prohibitive on the large dataset (*N*=561), failing to converge within five hours. PIMMS performed second best on the large dataset, but its accuracy dropped sharply when the MNAR proportion increased, and it exhibited a large performance gap between small and large sample sizes. By contrast, msBayesImpute maintained stable accuracy across both scenarios (Figure 3C).

To further dissect error sources, proteins were ranked into ten abundance groups, and over– and underestimation errors were quantified (Figure 3D). Overestimation was most pronounced among low-abundance proteins, where msBayesImpute showed marked improvements over PIMMS across all groups. Although msBayesImpute exhibited slightly higher underestimation than PIMMS on average, the difference was minimal.

Together, these results show that msBayesImpute delivers consistently strong performance on semi-synthetic HeLa proteomics data across varying MNAR proportions and sample sizes, underscoring its robustness and suitability for diverse experimental settings.

### Validation of msBayesImpute using serial dilution experiments on primary lung cancer tissues

Most existing MVI tools for mass-spectrometry proteomics have not been tested on truly experimental benchmarking data. This is largely because the ground truth for missing values is unknown, and any synthetic introduction of MNAR values inevitably relies on assumptions that may bias the evaluation toward certain methods. To minimize such bias, we employed serial dilution experiments to generate missing values with experimentally defined ground truth. Specifically, matched tumor and tumor-free samples from 10 lung adenocarcinoma patients were prepared in three dilution fractions: 100% (undiluted), 75%, and 50% of the original input concentration. Proteomic data from patient tissue samples are a particularly challenging matrix, often containing many missing values, making reliable information retrieval especially important. Values observed in the undiluted samples but absent in the diluted samples could therefore be regarded as ground truth. The experimentally generated missing values displayed an inverse relationship between protein abundance and dropout probability, consistent with a sigmoid-shaped dropout curve and in line with the assumptions validated in our synthetic and semi-synthetic benchmarking (Figure 4A–B).

**Figure 4.**
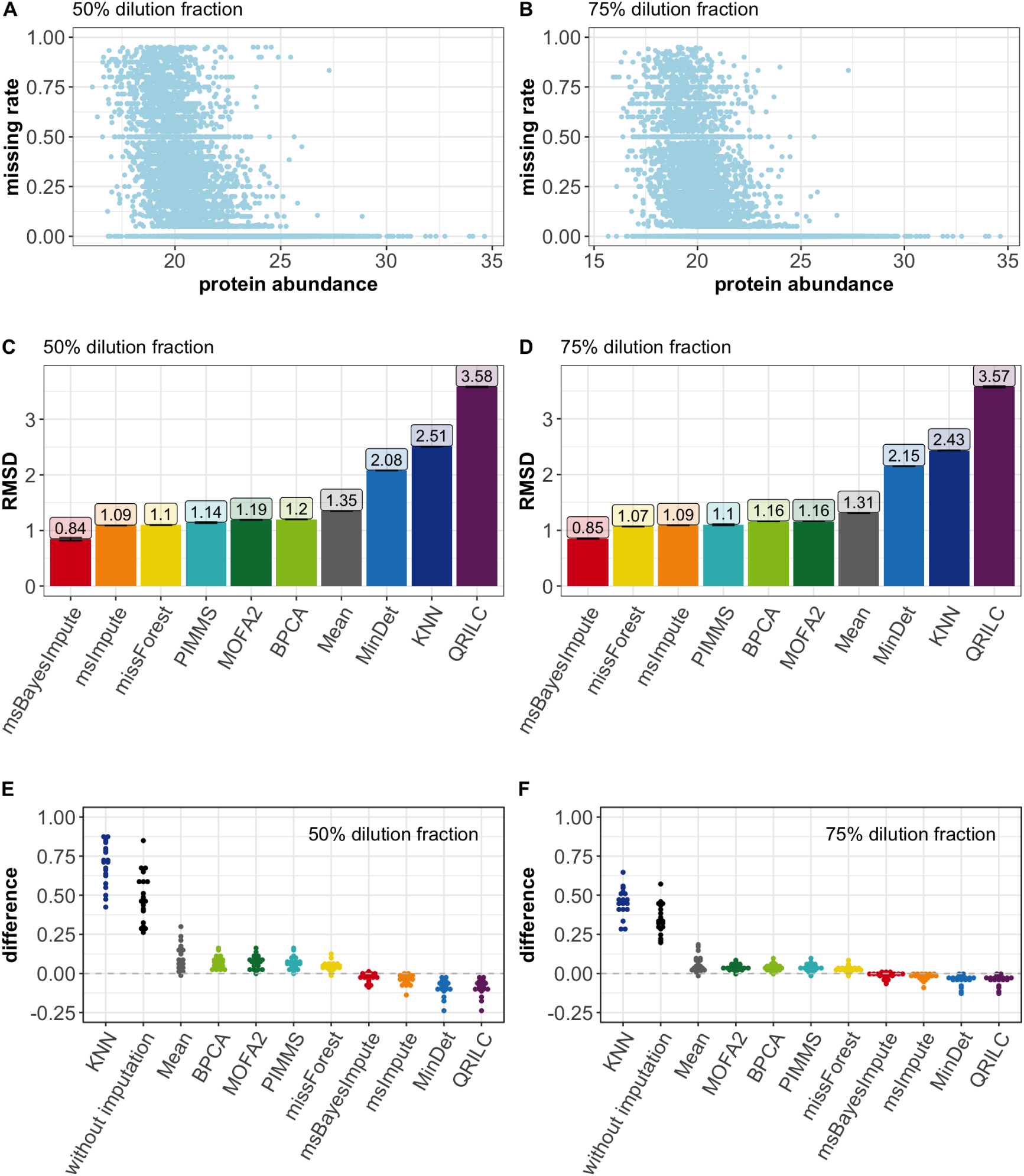
Benchmarking msBayesImpute using serial dilution experiments. (**A, B**) Inverse relationship between protein-specific missing rate and average protein abundance of the ground truth values observed in the dilution datasets. (**C, D**) Reconstruction accuracy of nine imputation methods plus MsImpute, evaluated on unbiased validation data generated from serial dilutions of the original sample concentration (75% and 50% relative to undiluted controls). RMSD between imputed values and ground truth values from the 100% (undiluted) dataset was used as the performance metric. (**E, F**) Errors in estimating sample-wise medians, calculated by comparing values from the original undiluted dataset (100%) with those from imputed or incomplete datasets at 75% and 50% fractions.

We next compared msBayesImpute with nine other MVI methods, including MsImpute, which models MAR/MNAR mixtures but is restricted to two-group experimental designs (e.g., tumor vs. tumor-free). Based on RMSD between imputed and ground truth values, msBayesImpute achieved the highest reconstruction accuracy, reducing error by ∼20% compared to the second-best method—MsImpute in the 50% dataset and missForest in the 75% dataset (Figure 4C–D). The performance ranking of other methods was consistent with the semi-synthetic HeLa benchmarks, with MinDet, KNN, and QRILC performing worst.

We further assessed the ability of msBayesImpute to recover true sample-wise overall intensity, represented by sample median abundance, which is essential for accurate normalization. As shown in Figure 4E–F, medians estimated from msBayesImpute-imputed matrices closely matched the ground truth and outperformed those from nine other methods or direct estimation on incomplete data. MsImpute yielded comparably accurate normalization, whereas KNN and unimputed data performed the poorest.

### Reliable detection of differentially expressed proteins with msBayesImpute

A central task of MS-based proteomics in biological and biomedical studies is to identify proteins whose abundances differ significantly between conditions or patient groups. However, MNAR missing values can hinder this process by reducing statistical power and introducing estimation bias, for example, by shifting group-wise means. To assess whether msBayesImpute improves differential expression (DE) analysis, we compared it with other MVI methods using the lung cancer serial dilution dataset. Because the true set of biologically differentially expressed proteins between tumor and tumor-free samples is unknown, we treated the DE proteins identified in the undiluted dataset at varying false discovery rate (FDR) thresholds as the ground truth. We then assessed the observed FDR and true positive rate (TPR) for each method relative to this reference. For comparison, we also included proDA, which bypasses imputation and performs DE testing directly on unimputed data by incorporating a probabilistic dropout model.

In the 50% dilution dataset, only msBayesImpute, KNN, and proDA successfully controlled the FDR at 5%, while all methods achieved control at 10% (Figure 5A). In the 75% dilution dataset, where fewer values were missing, all methods except mean imputation controlled the FDR at both 5% and 10% (Figure 5B). Across both datasets and across varying cutoffs, msBayesImpute consistently exhibited among the lowest actual FDR values, demonstrating stringent control in hypothesis testing. Importantly, despite this stringency, msBayesImpute achieved the highest TPR of all methods (Figure 5C–D).

**Figure 5.**
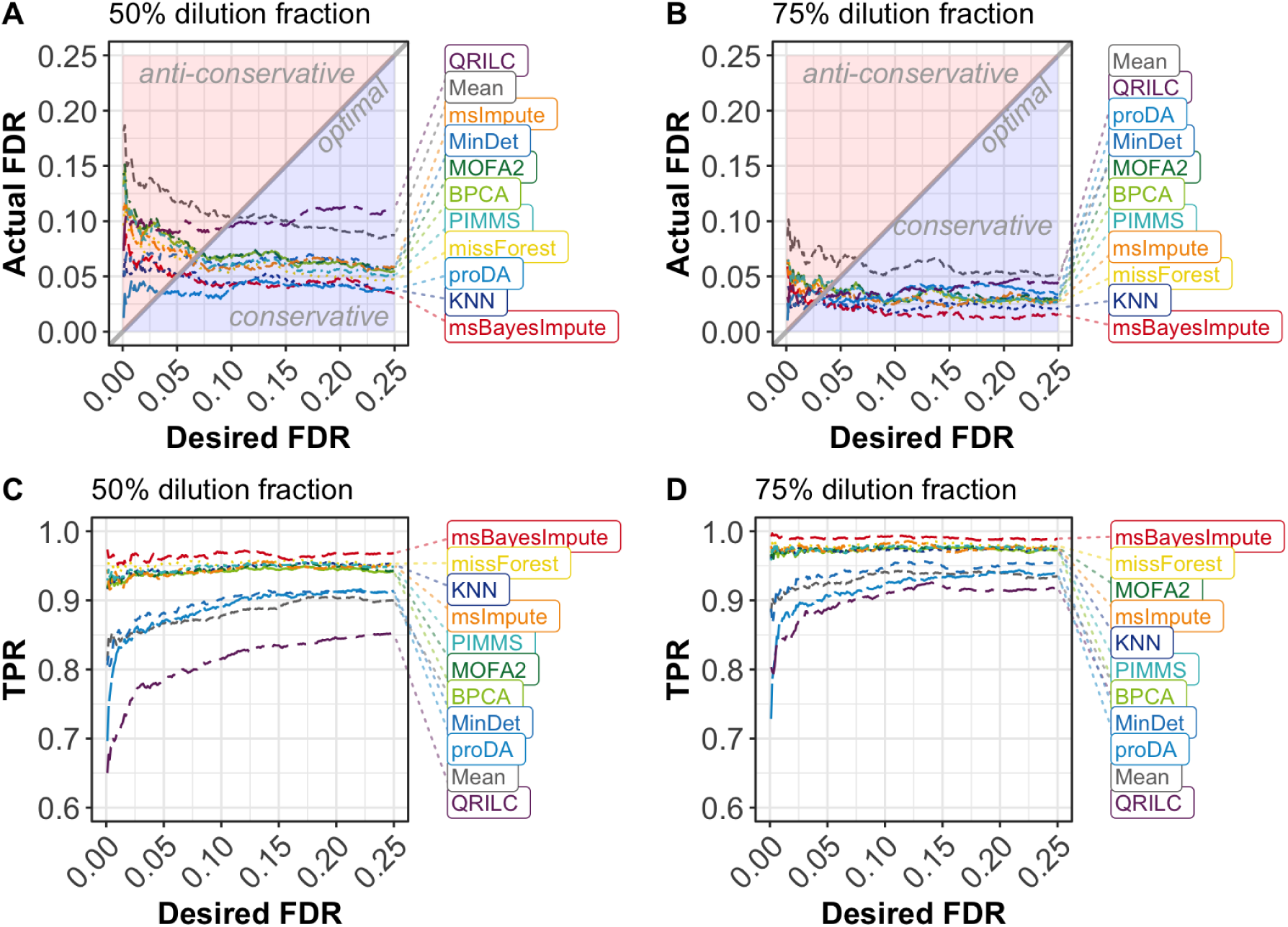
Performance comparison of differential expression analysis. (**A, B**) Relationship between desired and actual false discovery rate (FDR) in the 50% and 75% dilution datasets. Note that, due to the lack of a truly differentially expressed protein list, the desired FDR corresponds to the cutoff used to define differentially expressed proteins in the undiluted dataset (ground truth), while the actual FDR reflects the proportion of false discoveries when applying the same cutoff to imputed datasets. Therefore, the “ground truth” protein list will also grow when the desired FDR increases, which leads to a flatter look of the curves. A gray diagonal line indicates the optimal value. Light blue color indicates a conservative FDR, whereas orange indicates an anti-conservative FDR. **(C, D)** True positive rate (TPR) in the 50% and 75% dilution datasets, calculated as the proportion of ground truth positives (from the undiluted dataset) recovered in the imputed datasets.

We further quantified performance using the F1 score at a target FDR of 5% (Table 1). msBayesImpute achieved the highest F1 scores in both dilution datasets (0.955 and 0.982), followed by KNN (0.944 and 0.975) and MsImpute (0.931 and 0.970). ProDA yielded lower scores (0.914 and 0.937), and missForest, PIMMS, BPCA, and MOFA produced intermediate results. MinDet performed similarly to proDA, while QRILC and mean imputation failed to simultaneously control FDR and TPR, resulting in the poorest performance.

**Table 1.** F1 scores measurements from the Lung cancer serial dilution experiments. The F1 score, reflecting the balance between true positives and true negatives, was calculated with the desired false discovery rate (FDR) fixed at 0.05. Ten imputation methods, along with proDA, were evaluated.

| Experiment | Method | F1 score |
| --- | --- | --- |
| 50% dilution experiment dataset | msBayesImpute | 0.955 |
|  | KNN | 0.944 |
|  | missForest | 0.942 |
|  | PIMMS | 0.931 |
|  | msImpute | 0.931 |
|  | BPCA | 0.928 |
|  | MOFA2 | 0.928 |
|  | proDA | 0.914 |
|  | MinDet | 0.906 |
|  | Mean | 0.875 |
|  | QRILC | 0.842 |
| 75% dilution experiment dataset | msBayesImpute | 0.982 |
|  | KNN | 0.975 |
|  | msImpute | 0.970 |
|  | missForest | 0.970 |
|  | MOFA2 | 0.968 |
|  | PIMMS | 0.968 |
|  | BPCA | 0.966 |
|  | MinDet | 0.956 |
|  | proDA | 0.937 |
|  | Mean | 0.928 |
|  | QRILC | 0.924 |

Together, these results demonstrate that msBayesImpute most effectively balances sensitivity and specificity, enabling reliable detection of differentially expressed proteins and recovery of robust biological signals.

## Discussion

Mass spectrometry–based proteomics is now indispensable for advancing our knowledge of biological processes and their dysregulation in disease, yet missing values remain a persistent challenge that can compromise downstream analyses. Robust and accurate imputation is therefore critical. Here, we present msBayesImpute, a Bayesian matrix factorization–based method with a probabilistic dropout model that addresses both MAR and MNAR missingness in a data-driven way.

Our benchmarks across synthetic, semi-synthetic, and experimental datasets demonstrate that msBayesImpute offers clear advantages over existing MVI tools. It achieves higher imputation accuracy, particularly for low-abundance proteins that are most affected by MNAR dropout. Unlike many conventional methods, it also improves the recovery of sample-wise intensities (e.g., median abundance), enabling more accurate normalization and avoiding systematic biases that can propagate into downstream analyses. Moreover, msBayesImpute consistently enhanced the detection of differentially expressed proteins, showing both stringent FDR control and superior sensitivity compared with other MVI methods and proDA.

Beyond its performance, msBayesImpute is characterized by flexibility and ease of use. It requires no parameter tuning and learns dropout curves directly from the data, without the need for predefined experimental designs or preprocessing such as filtering or normalization. This distinguishes it from methods like MsImpute (Hediyeh-Zadeh, Webb and Davis, 2023) and proDA (Ahlmann-Eltze and Anders, 2020), which are restricted to specific designs (e.g., two-group comparisons). As a result, msBayesImpute is broadly applicable, ranging from exploratory data analysis to machine learning pipelines and signaling network modeling. It performs robustly on both small and large datasets, avoids the heavy sample size requirements of deep learning approaches such as PIMMS (Webel *et al*., 2024), and is substantially more computationally efficient than BPCA and missForest. To facilitate adoption, we provide implementations in R and Python, along with a Shiny app and Docker image for non-programmers.

Despite these strengths, msBayesImpute has some limitations. Its use of a variational Bayesian framework means that uncertainty quantification is generally underestimated (Blei, Kucukelbir and McAuliffe, 2017), which could limit its integration into downstream statistical inference or modelling. Extending the model to provide uncertainty estimates would improve control of false positives and negatives and increase reproducibility. Another limitation of msBayesImpute is its reduced performance on extremely small datasets (e.g., fewer than 10 samples). In such cases, robust factorization and reliable estimation of protein-specific dropout curves become difficult in a purely data-driven manner. A potential solution would be to develop a pre-trained empirical model based on the growing number of large-scale proteomics datasets available in public repositories. Such a framework could provide informative priors to guide imputation in very small studies, thereby extending the applicability of msBayesImpute. Finally, although msBayesImpute is substantially more efficient than computationally heavy methods such as BPCA and missForest, it remains slower than simpler approaches (e.g., MinDet, QRILC) and other factorization-based methods (e.g., PIMMS, MOFA) (Figure 2E and Supplementary Figure 2). The main computational bottleneck is the inference of protein-specific dropout curves, which could be addressed through the use of pre-trained empirical models, more efficient computational engineering, and implementation in faster backends (e.g., C++ libraries instead of Python).

Several promising directions remain for extending and applying msBayesImpute. First, msBayesImpute can be applied to single-cell proteomics, where dropout is pervasive and largely MNAR, and could address a pressing challenge in this emerging and exciting field. Second, its probabilistic dropout model can be adapted to other MS-based data types, including phosphoproteomics, metabolomics, and lipidomics, broadening its utility. In addition, integrating msBayesImpute with multi-omics factor analysis (MOFA) represents an exciting technical opportunity. While MOFA effectively models MAR missingness in multi-omics data, it does not explicitly account for MNAR patterns. Because msBayesImpute and MOFA share similar Bayesian factorization frameworks, it should be technically feasible to combine them, allowing users to optionally enable dropout modeling for MS-based datasets and thereby improve integration performance. Finally, incorporating more flexible dropout models, such as zero-inflated negative binomial functions, could extend its applicability to transcriptomics and beyond.

In conclusion, msBayesImpute provides a robust, versatile, and user-friendly framework for handling missing values in MS-based proteomics. By improving imputation accuracy, normalization, and the detection of biological signals, it contributes to more reliable and reproducible proteomics research and opens avenues for application across diverse omics domains.

## Methods

### MsBayesImpute structure

The msBayesImpute model integrates Bayesian matrix factorization with a probabilistic dropout mechanism to impute missing values in numeric matrices representing protein abundances across samples measured by mass spectrometry. Starting from MS-based data *X* ∈ *R^N^*^×*D*^with missing values, where N is the number of samples and D is the number of features in the data matrix, msBayesImpute decomposes the original data matrix into lower-dimensional latent factor and weight matrices,

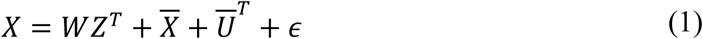

Where *W* denotes the weight matrix, and *Z* denotes the factor matrix. 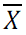 is a vector with the length of D, representing the mean value (intercept) per feature. Due to the presence of missing values, the mean or median abundance of all features in a sample, which is important for sample normalization, could not be accurately calculated by only taking the observed values. Therefore, 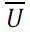 is introduced as a vector of the length of N to model the true difference of mean abundance across samples. The residual noise matrix is represented by ϵ. *W*, *Z*, 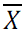 and 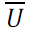 are the parameters to be estimated with the variational Bayesian approach. After the inference of *W*, *Z*, 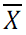 and 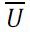 the complete data matrix without missing values can be reconstructed with formula (1). If the missing data points are MAR, they can be simply ignored during the inference process, which is how Bayesian PCA or MOFA performs missing value imputation. However, in mass-spectrometry data, the missing values were usually not generated at random. Proteins with lower abundance tend to contain more missing data points. This trend can be described using a probabilistic model introduced by Ahlmann-Eltze et al. (Ahlmann-Eltze and Anders, 2020), revealing that the likelihood of missing value decreases as protein abundance increases.

The probabilistic dropout model 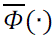 is mathematically represented by the cumulative distribution function of a Gaussian distribution Φ(·) and can be incorporated into the likelihood of a Bayesian factorization model:

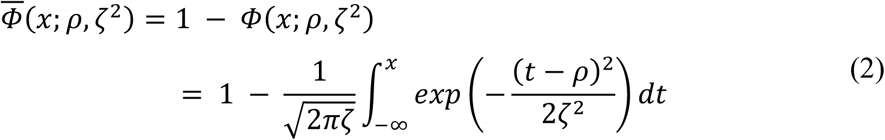

### Likelihood models

The Bayesian matrix factorization framework of msBayesImpute integrates two likelihood models: the Gaussian distribution for modelling observed values and the probabilistic dropout distribution for modelling missing patterns. The probabilistic dropout distribution, 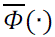, is modelled as the cumulative density function of a Gaussian distribution, forming a sigmoid-shaped curve characterized by two key parameters, i.e., the inflection point ρ and the scale ζ. Specifically, we established feature-wise probabilistic dropout distributions to model the missing likelihood, accounting for the varying extent of missing patterns associated with each protein. All data points *x_dn_*_’_, both missing and observed, carry missingness information and are therefore fitted using probabilistic dropout models:

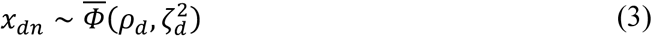

On the other hand, the observed data follow Gaussian distributions δ(·) at the same time:

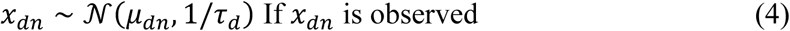

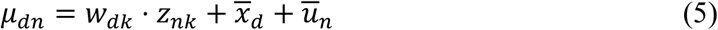

Where μ*_dn_* corresponds to the reconstructed values using a list of factorization parameters, including 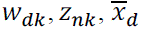 and 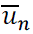, and 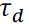 stands for precision per feature.

### Probabilistic dropout priors

We assume the missing information can be inferred from the given data *X*, with the dropout patterns varying slightly across features. For the inflection point of each dropout mode, they are assumed to share the global inflection point ρ_0_ derived from *X*,

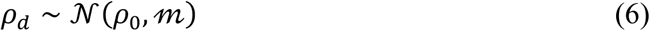

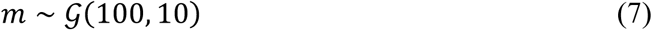

Where *g*(·) represents a gamma distribution. For the scale of each dropout mode, we assume they do not directly stem from the global scale of ζ_0_, but rather from hierarchical priors, reflecting the high heterogeneity in scales as compared to the inflections.

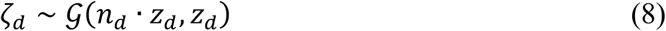

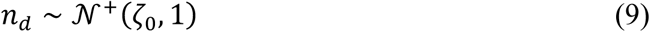

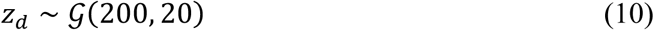

Where *n_d_* ∈ *R*^+^ indicates the constant negative relationship between MS-based abundance and missing rates to represent the missing value patterns observed in MS proteomics data. The global ρ_0_ and ζ_0_ can be directly achieved from the given data *X*,

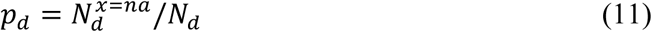

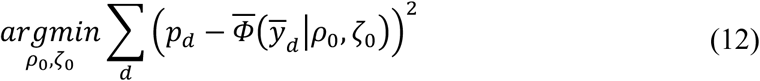

Here, *N_d_* corresponds to the number of samples and 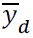 denotes imputed values by MinDet method.

Moreover, the hyperpriors for the variance of inflection and the rate of scale, i.e., *m* and *Z_d_*, can influence performance as protein-specific missing patterns are assumed to be heterogeneous. We benchmarked these hyperpriors using both synthetic and semi-synthetic datasets through a grid search. The combinations of (100, 10) for the gamma distribution of the variance *m* and (200, 10) for the rate *Z_d_*, indicated in formulas (7) and (10) led to the best and most stable performance.

### Matrix factorization priors

The parameters in formulas (4) and (5) are inferred from a list of priors. The precision τ*_d_* of the Gaussian likelihood can be modelled using the following Gamma and Gaussian hyperpriors,

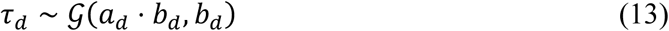

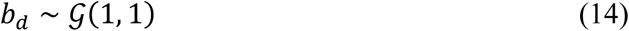

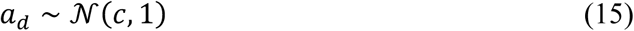

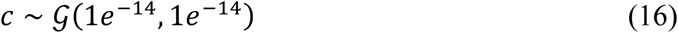

Among 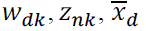 and 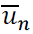, Gaussian priors were assigned to 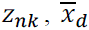 and 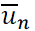, while a shrinkage prior was employed for *w_dk_* to reduce spurious correlations and false positives. We provided the Horseshoe prior for feature-wise sparsity:

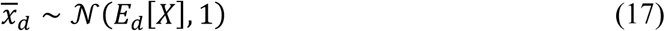

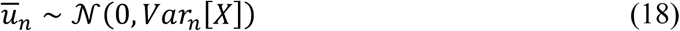

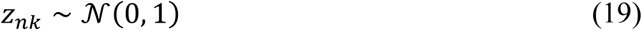

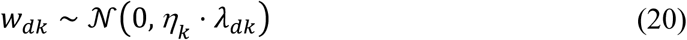

Here *k* ∈ {1, …, *K*} are latent factors, *E_d_* stands for empirical mean and *Var_n_* is empirical variance. *η_κ_* are factor-specific variables used to estimate the general level of activeness for each factor, while λ*_dk_* is a feature-specific shrinkage parameter allowing each element in W to escape the factor-specific sparsity. λ*_dk_* was sampled from a positive Half-Cauchy distribution *C*(⋅), and *η_κ_* was sampled from a Beta distribution. To automatically learn *η_κ_*, and λ*_dk_* we used uninformative fixed priors,

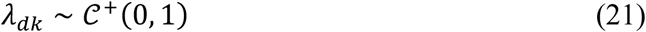

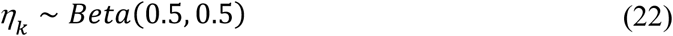

Finally, the joint probability density function is generated by these prior and likelihood distributions:

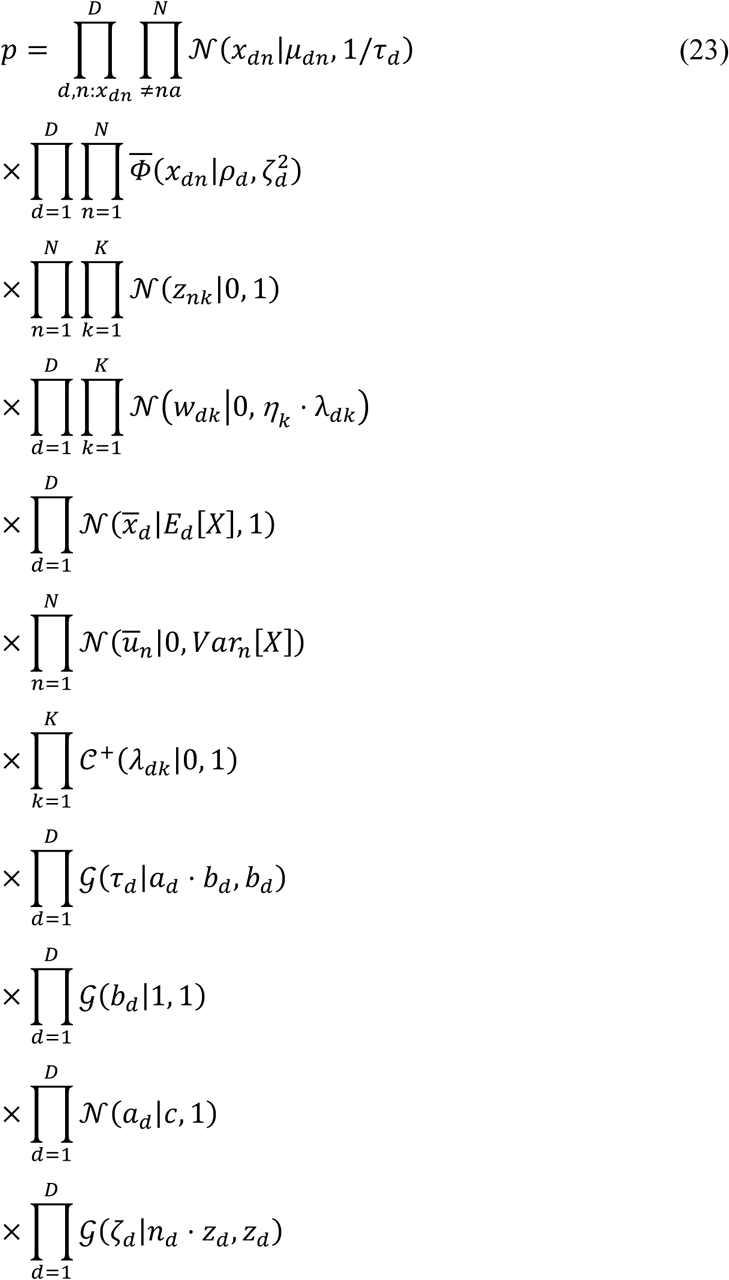

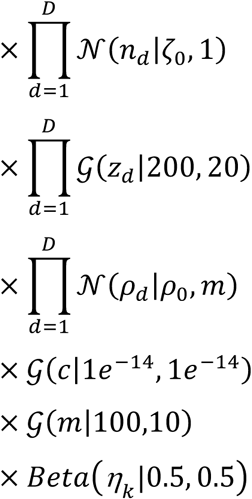

### Parameter inference

The msBayesImpute is a matrix factorization-based model using a probabilistic Bayesian framework and stochastic variational inference (SVI). The main idea is to approximate the intractable posteriors by maximizing the evidence lower bound (ELBO). The msBayesImpute employs Adam, a stochastic gradient descent optimizer with a learning rate of 0.05 and also employs a Normal distribution with a diagonal covariance matrix as an automatic guide to approximate the posterior distribution. The convergence includes two modes, slow and fast, with training terminating when the supervised loss difference remains below a specified threshold associated with the convergence mode for three consecutive iterations. The msBayesImpute model consists of two modules, the initialization block and the refinement block. The maximum number of training iterations per block is 30,000.

There are mainly two initialization steps: calculating the global ρ_0_ and ζ_0_ given data X with its imputed values using MinDet, indicated in formulas (11) and (12), and searching for significant latent factors. The model can automatically learn the number of latent factors by filtering out insignificant factors under a threshold of explained variance of 1%. Alternatively, the user can set the factor number or the explained variance threshold during model initialization. In the former case, the threshold is ignored. A minimum of one latent factor is retained. Additionally, if variances differ significantly across features, but the sample size is too small for the model to capture this variation, an additional initialization step is applied. msBayesImpute can estimate an initial global residual to replace the hyperparameter *c* in formula (16) as an informative prior for precision τ, by performing principal component analysis (PCA) on imputed values from MinDet.

To further refine the model for more accurate imputation after initialisation, msBayesImpute alternates between estimating the matrix factorization parameters and probabilistic dropout parameters. When estimating one set of parameters, the other set of parameters are fixed. The refinement iterations consist of an initial factorization optimization, followed by a dropout optimization, and conclude with a second factorization optimization. The same termination criterion is applied in each optimization as in the initialization steps, and the specific epoch per optimization is 10,000.

This model is established in Python using the probabilistic programming language Pyro. An R wrapper package has been implemented for R users, and an RShiny platform as an interactive interface has been established for non-programmers.

### Setup of other imputation methods

The other MVI methods that were benchmarked against msBayesImpute include PIMMS, MissForest, MOFA, BPCA, MsImpute, KNN, MinDet, QRILC, and ProDA, with feature-wise mean used as the baseline method. All methods, except for PIMMS, which was only available in Python, were accessible in R.

The PIMMS has three approaches within the pimmslearn Python package (Webel *et al*., 2024), including collaborative filtering (CF), denoising autoencoder (DAE), and variational autoencoder (VAE). Among these, DAE focuses on values reconstruction and therefore was used for benchmarking. MOFA imputation was performed using the MOFA2 package (R version 1.14.0) (Argelaguet *et al*., 2018, 2020; Velten *et al*., 2020) with the Automatic Relevance Determination (ARD) prior on the weights. BPCA was conducted using the pcaMethods package (R version 1.96.0) with centring enabled. For factor analysis-based models, we initialized msBayesImpute and MOFA with automatic factor activation, setting the explained variance at 1%, while setting the number of BPCA factors between the pruned factors of msBayesImpute and MOFA. KNN, MinDet and QRILC imputations were carried out using the MsCoreUtils package (R version 1.16.1) (Rainer *et al*., 2022). By default, the number of neighbours in KNN is set to 10, and the maximum missing percent in columns is 80% by default in the dilution experiment, 85% in the semi-simulated data, and 85% –95% in simulated data, while the quantile parameter in MinDet is set to 0.01. MissForest was run using the missForest package (R version 1.5) (Stekhoven and Bühlmann, 2012, 2022), which doesn’t require setting parameters, although the minimum number of observed values for each feature is 2, while MsImpute was executed with the msImpute package (R version 1.14.0), requiring a minimum of 4 observed values per feature. Lastly, ProDA was executed with default settings using the proDA package (R version 1.18.0).

### Data simulation

To evaluate whether the msBayesImpute model could capture the ground truth parameters, we first simulated data from the generative model msBayesImpute was based on with varying numbers of features and factors. For each simulated dataset without missing values, a weight matrix and a factor matrix with specific dimensions were drawn from a standard Gaussian distribution. After the multiplication of weight and factor matrices, intercepts for each feature, sampled from Gaussian distributions parameterized by specified means and variances, were added. To mimic the variability observed in real proteomics, Gaussian noise was also introduced, with precision parameters per feature drawn randomly from a Gamma distribution.

To introduce missing values that resemble the real-world missing pattern in label-free MS proteomics, two methods were used. One method generates missing values with the probabilistic dropout function msBayesImpute based on, with individual inflection and scale parameters per feature drawn from a Gaussian distribution with a chosen location and standard deviation. This method is useful for validating our model but may introduce a favourable bias towards our model when benchmarked against other MVI models. On the other hand, this kind of probabilistic dropout likelihood is currently only suitable for label-free methods rather than others like TMT-or SILAC labelled MS data. Therefore, another method, which was introduced by Lazar et al. (Lazar *et al*., 2016), generates missing values based on mixing two missing type mechanisms, i.e., MAR and MNAR, with a manual specific of mixture proportions. The MAR mechanism creates missing values randomly in the quantitative dataset. For the MNAR mechanism, a threshold matrix with the same dimension of the original dataset is sampled from a Gaussian distribution, centred at a chosen quantile of the original data and with a standard deviation set to 0.3 times that of the original distribution. Values in the original dataset are designated as missing if they fall below their corresponding thresholds. A key strength of this method is its precise control over the missing rate associated with each type of missingness and to avoid bias during benchmarking.

To examine the effect of missing rates and sample sizes on prediction across different imputation methods, datasets were systematically generated with missingness rates ranging from 10%, 30% to 50% and sample sizes from 20, 100 to 200, using the probabilistic dropout function as missing rate and sample sizes are critical factors influencing the performance of imputation methods (Jin *et al*., 2021).

### Generation of the semi-synthetic benchmarking dataset

For benchmarking, a semi-synthetic dataset, where artificially generated missing values were introduced into a label-free MS-based proteomics data derived from HeLa cell lines (ProteomeXchange ID: PXD042233). This dataset, which consists of two protein expression matrices with a sample size of 564 and 50, respectively, was used as the benchmark dataset for the PIMMS method, as sample size plays a significant role in influencing performance. The missing value generation mechanism that mixes MAR and MNAR with a fixed proportion by Lazar et al. (Lazar *et al*., 2016), which has been used for benchmarking PIMMS (Webel *et al*., 2024) and by others in similar benchmarking studies (Jin *et al*., 2021; Harris *et al*., 2023;Wei *et al*., 2018), was used to introduce missing values in the semi-synthetic data for a fair comparison among different MVI methods. In addition, this missing generation approach can control MNAR rates, which can be employed to assess the robustness of imputation methods when the composition of the missing pattern changes.

### Generation of the benchmarking dataset with serial dilution experiment

To better recapture the missing value pattern in a real-world situation, the missing values were experimentally introduced through mass-spectrometry measurements of 20 paired tumour/tumour-free tissues from 10 lung cancer patients without dilution and with dilution fractions of 75% and 50%.

Human lung tissue samples were obtained from patients with lung adenocarcinoma and provided by the Lung Biobank Heidelberg, a member of the accredited Tissue Bank of the National Center for Tumor Diseases (NCT) Heidelberg. The study was approved by the Ethics Committee of the Medical Faculty of the University of Heidelberg (approval number: S-270/2001 (biobank vote) and S-154/2018 (study vote). Written informed consent was obtained from all patients. Experienced pathologists oversaw tumor dissection, selecting tumor enriched regions as well as corresponding tumor free areas. Samples were received snap-frozen and were kept on dry ice during processing. For sample preparation two slices each were transferred to separate tubes containing a small spoon of protein extraction beads (Protein Extraction Beads, diagenode #C20000021) for tissue lysis. Tissue was stored at –80 °C. After thawing on ice, between 50 μL to 150 μL of 1x RIPA lysis buffer with 1% NP40 and 0.1% SDS was added (1% NP40, 0.1% SDS, 1% sodium deoxycholate, 25 mM Tris-HCl pH 7.6, 150 mM NaCl, 10 mM NaF, 1 mM Na3VO4, 250 U/mL benzonase, 10 U/mL DNase, 0.1% AEBSF, 0.1% aprotinin), depending on the size of tissue slices. To ensure a full coverage with lysis buffer, samples were spun down 1 min at 14 000 rpm at 4°C prior to incubation at 60 °C for 2 h. Samples were centrifuged for 1 min at 14 000 rpm at 4°C. Samples were sonicated in a Bioruptor Pico (Diagenode) for 10 cycles (30 s on /30 s off, easy mode, 4 °C). After an additional centrifugation step (14 000 rpm, 10 min, 4 °C) the supernatant was collected and the concentration of tissue lysates were determined by using the BCA Assay Kit (Thermo Scientific Pierce # A55864).

Protein reduction, alkylation, cleanup and tryptic digestion were performed by following SP3 protocol on an automated liquid handling platform (Agilent Bravo). 55 μg protein was used for further procedure and prepared in a 96 well plate (Superplate, Thermo #AB-2800) in a total volume of 30 μL of 100 mM triethylammonium bicarbonate (TEAB, Sigma Aldrich #T7408-100ml). Prior to running the protocol on Agilent Bravo, 40 mM tris-(2-carboxyethyl)-phosphine (TCEP, Sigma Aldrich # C4706-2G) and 160 mM 2-chloroacetamide (CAA, Sigma Aldrich # C0267-100G) were prepared as stock solutions for reduction of disulfide bonds and alkylation. For sample cleanup, master mixes of Sera-Mag Speed Beads A (hydrophobic coating) and B (hydrophilic coating) (GE Healthcare, Beads A #65152105050250, Beads B # 45152105050250) were prepared in a ratio of 1:1 by using 100 μL of each bead suspension. To remove the acidified solution in which the beads are stored in, the bead master mix was cleaned by placing the tubes on a magnetic rack until all beads were located to the magnet (approx. 1 min) and the solution appeared clear. Supernatant was removed, beads were resuspended in 200 μL MS-compatible H2O and put back to the magnetic rack. This procedure was performed in total three times. Beads of each master mix were resuspended in a volume of 90 μL to achieve a final volume of 100 μL. To enhance protein binding to the beads surface, 100 % MS-compatible ethanol was prepared as well as 80% MS-compatible ethanol for protein cleanup procedure. For protein digestion buffer a solution containing 2.2 μg Trypsin Gold (PROMEGA # V5280) / 75 μL 100 mM TEAB was prepared (1:25 protease to protein ratio). After preparation of the deck overlay on Agilent Bravo, the following steps were executed. Reduction and alkylation were performed by adding 15 μL of TCEP (final conc. 10 mM) and CAA (final conc. 40 mM) followed by an incubation step at 95 °C for 5 min. Protein cleanup was initiated by the addition of 5 μL beads master mix and 65 μL 100 % MS-compatible ethanol to a final concentration of 50 % (v/v). The sample plate was shaking for 15 min at 800 rpm to allow sufficient protein binding. After incubation, the sample plate was moved to a magnetic rack, supernatant was removed after 1 min and beads were washed by resuspending in 80% MS-compatible ethanol. Wash steps were executed three times. As a final step 75 μL of digestion buffer was added to the beads and were incubated 16 h at 37 °C on a heating block of Agilent Bravo. To avoid sample evaporation, the plate was sealed with sealing foil as soon as tryptic digestion started. After incubation, the sample plate was centrifuged for 1 min at 14 000 rpm. Sealing foil was removed and digested proteins were transferred to a new plate using the magnet and 96LT pipetting head on Agilent Bravo deck. After lysate collection peptides were dried down by vacuum centrifugation at 45°C. Samples were reconstituted in 130 μL MS-compatible 80 % ACN + 0.1 % FA. Afterwards, 5 μg (equal to 12.3 μL) were transferred to a new 96 well plate and dried down by vacuum centrifugation at 45°C for full proteome measurement and dilution series. The remaining 50 μg were used for a different procedure. Finally, the sample plate was stored at –20 °C until reconstitution for MS measurement. For reconstitution and dilution, loading buffer (0.1% formic acid (FA), 2% ACN in MS-compatible H2O) was used. Digested, undiluted peptide samples (100%) were dissolved in 15 μL loading buffer. Out of this initial reconstitution, a 75% dilution was prepared, followed by the 50% dilution based on the previous generated dilution (75%).

Nanoflow LC-MS/MS was performed by coupling an UltiMate 3000 to an Orbitrap Eclipse Tribrid mass spectrometer (Thermo Scientific). Reconstituted and diluted samples were spiked in by using an injection volume of 2 µL. Peptides were loaded to a trapping column (PEPMAP NEO C18 5 µm 0.3 mm x 5 mm, 1500 bar) at a flow rate of 30 µL/min in 100% buffer A (0.1% FA in MS-compatible H2O). After loading onto an analytical column (100 μm ID × 15 cm length, packed in-house with Reprosil-Pur 120 C18-AQ, 1.9 μm resin, Dr. Maisch) peptides were separated using a 85 min gradient from 2% to 38% of buffer B (0.1% FA, 80% ACN in MS-compatible H2O) at a 300 nL/min flow rate. The Orbitrap Eclipse™ Tribrid™ mass spectrometer was operated in data-independent mode (DIA), automatically switching between full scan MS and MS2. Full scan MS spectra were acquired in the Orbitrap at 120 000 resolution, m/z ranging from 350 to 1400, after accumulation to the set target value of 300% (100% = 1.2e6) with a maximum injection time of 45 ms. The DIA method covers 30 isolation windows with a dynamic range of 400 to 1000 m/z and uses a higher energy collisional dissociation (HCD) for fragmentation at normalized collision energy (N)CE of 28 %. MS2 spectra were acquired in the Orbitrap at a resolution of 30 000 with a target value of 1000 % (100 % = 5e5) charges after accumulation for a maximum of 54 ms. Generated data was measured in positive ion mode.

DIA-NN with default parameters was used to process the raw mass-spectrometry data from the diluted and undiluted samples to quantify protein abundance. The values of data points that were observed in the undiluted samples, while unobserved in the diluted samples, were used as the ground truth values for benchmarking.

### Data pre-processing

A standardized set of preprocessing steps was applied to the simulated, semi-synthetic and experimental datasets for all imputation methods to ensure comparability. Proteins that were missing in all samples after introducing missing values were first removed. Certain imputation methods, including MissForest and MsImpute, require a minimum of 2 and 4 observations per protein, respectively. For specific comparisons, such as with PIMMS, we followed the same processing steps outlined in their published papers to ensure comparable and reliable results, like removal of proteins with a missing rate greater than 75% and samples with a missing rate greater than 50%. For the semi-synthetic and experimental data, a log transformation of base 2 was applied to the spectra counts prior to imputation. Before comparing differential expression, we also converted the imputed data back to raw abundance expression in order to apply Variance Stabilization Normalisation (VSN), which was proven to outperform other commonly used normalisation methods in differential expression tasks, such as the median normalisation approach (Välikangas, Suomi and Elo, 2016).

### Performance evaluation

For benchmarking the accuracy of imputation, Root Mean Squared Deviation (RMSD) was used to compare the ground truth values and imputed values:

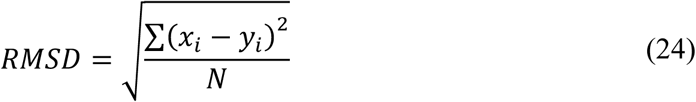

Where *x* and *y* stand for ground true and imputed values, respectively. Lower RMSD reflects greater imputation accuracy. All MVI methods were performed under different random seeds for 5 repetitions to obtain 95% confidence intervals and to assess the stability of each method.

Moreover, applying normalization before imputation may introduce bias, making it essential to assess how well different imputation methods handle subsequent normalisation compared to the original datasets with missing values as a baseline dataset. As an additional baseline, we also considered a prefiltered dataset in which proteins with more than 50% missing values were removed, since 50% filtering is a common analytical strategy. In this study, we assessed performance by comparing sample-wise median measurements from the imputed datasets to those from the baseline datasets.

To perform differential expression on ground truth and imputed expression matrices, linear models with Empirical Bayes moderated t-statistics implemented in the limma package (R version 3.60.6) (Ritchie *et al*., 2015) was used, except for ProDA, which performs differential expression analysis directly on an unimputed matrix. The measurements, including the true positive rate (TPR), false discovery rate (FDR), and F1 score, are used to evaluate the accuracy of a statistical test. In the undiluted expression data, proteins with adjusted p-values less than 0.05 were considered as actual positives, and those identified from the imputed matrices were treated as predicted positives. Predicted positives that overlapped with actual positives were classified as true positives (TP) and those that did not were considered false positives (FP), while proteins in actual positives rather than in predicted positives were designated as false negatives (FN). Therefore,

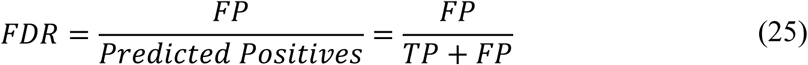

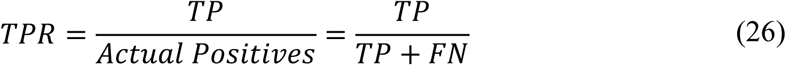

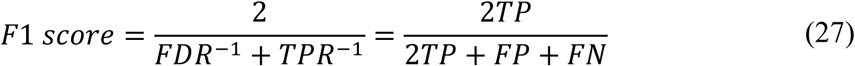

The F1 score, which harmonizes precision and recall, serves as an indicator of imputation performance, with higher values indicating improved accuracy.

## Data availability

The public HeLa cell lines data, consisting of 564 and 50 samples used in this paper, can be accessed at https://github.com/RasmussenLab/pimms. The processed proteomic dataset of the paried tumor and tumor-free tissue samples from 10 lung cancer patients is available at https://github.com/Lu-Group-UKHD/msBayesImpute_manuscript.

## Code availability

The code for msBayesImpute package can be found at: https://github.com/Lu-Group-UKHD/msBayesImpute. A workflowR project with the analysis code, figures, and results associated with the manuscript is available at https://github.com/Lu-Group-UKHD/msBayesImpute_manuscript.

## Author contributions

J.L. conceptualized and initiated the study. J.L. and B.V. guided the computational analysis. J.H. assembled the data, performed the analyses and wrote the software. M.A.S., L.V.K. and H.W. collected, prepared and provided the lung cancer patient samples. B.H., F.G., K.B., M.S. and U.K. performed the mass-spectrometry analysis of the lung cancer patient samples. J.L. and J.H. wrote the manuscript with contributions from all coauthors. All authors read and approved the final version of the manuscript.

## Acknowledgements

Junyan Lu and Jiaojiao He were supported by the SMART-CARE project funded by the Federal Ministry of Research, Technology and Space (BMBFTR) of Germany (funding code: 031L0213) and the Deutsche Forschungsgemeinschaft (DFG, German Research Foundation) – SFB 1709/1 2025 – 533056198. We thank Jana Braunger for providing technical support with Pyro programming.

