## Supplementary Figure 1 and 2 for "msBayesImpute: A Versatile Framework for Addressing Missing Values in Biomedical Mass Spectrometry Proteomics Data"

### Supplementary Figures

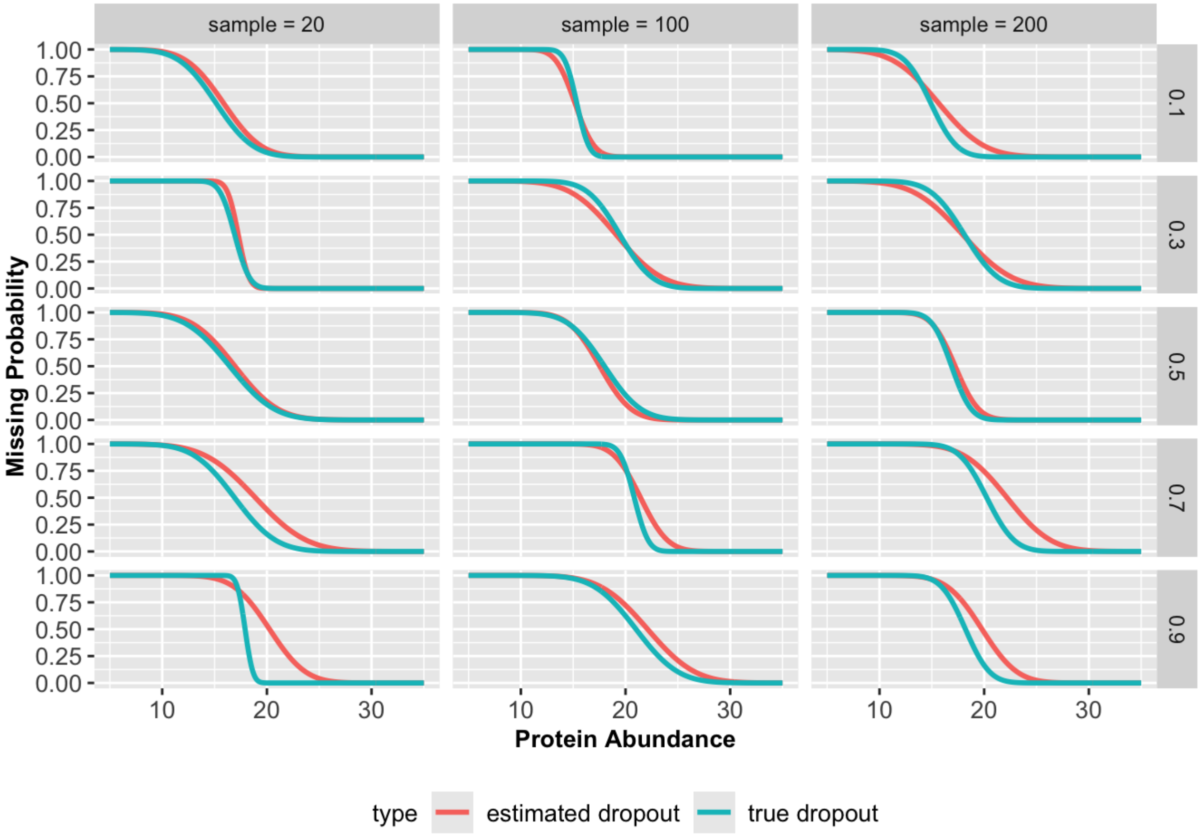


**Supplementary Figure 1: Prediction of dropout patterns.** Fifteen protein-specific dropout patterns were randomly selected under different missing rates and sample sizes.

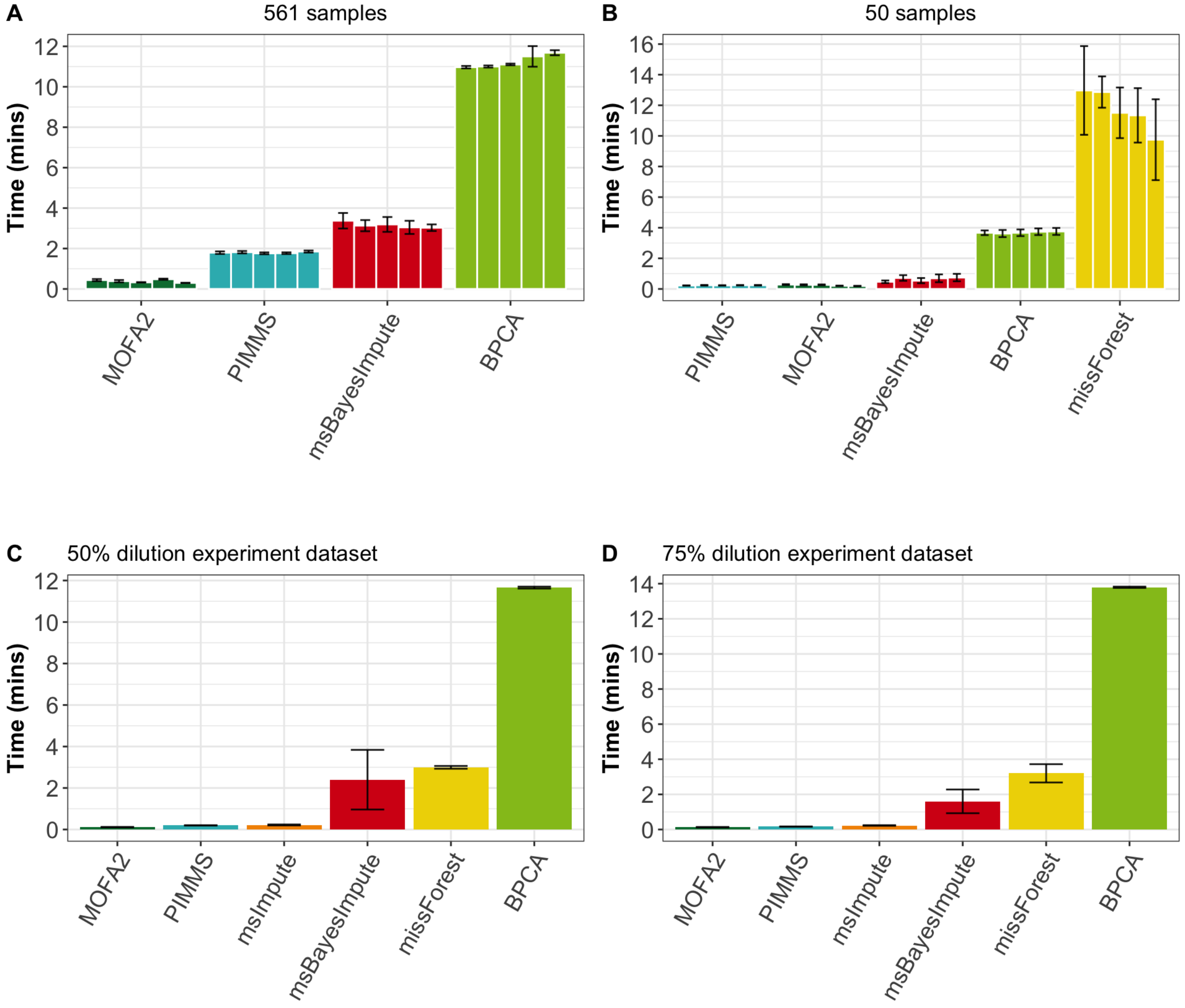


**Supplementary Figure 2. Computational time for imputation of proteomic datasets from HeLa cell line and lung cancer serial dilution experiments. (A)** Runtime comparison of MOFA2, PIMMS, msBayesImpute, and BPCA. For large-sample datasets, msBayesImpute required ~3 minutes, whereas simpler methods such as protein-wise mean, KNN, MinDet, and QRILC completed in <1 second. **(B)** Performance under small-sample datasets: msBayesImpute completed within ~1 minute, while missForest required ~10 minutes. **(C–D)** Inclusion of msImpute in the lung cancer serial dilution datasets: msBayesImpute required ~2 minutes, whereas BPCA showed the highest computational cost among the evaluated methods.
